# Dog-assisted scat sampling reveals genetic diversity, structure, and relatedness in black-footed cats (*Felis nigripes*)

**DOI:** 10.64898/2026.11.09.765305

**Authors:** Michelle M. Schroeder, Alexander Sliwa, Byron Weckworth, Beryl Wilson, Martina Küsters, Victoria Grant, Molly Schumer, Nadine Lamberski, Imogene Cancellare, Kristine Pilgrim, Greg Distiller, Jacqueline M. Bishop

**Affiliations:** Institute for Communities and Wildlife in Africa, Department of Biological Sciences, University of Cape Town, South Africa; Cologne Zoo, Cologne, Germany; Black-footed Cat Working Group: Black-footed Cat Research Project Namibia, Namibia; McGregor Museum, Zoology Department, Kimberley, South Africa; Conservation Genetics Program, Panthera, New York, New York, USA; Mountain Lion Foundation, Concord, California, USA; San Diego Zoo Wildlife Alliance Escondido, California, USA; Statistics in Ecology, Environment and Conservation (SEEC), Department of Statistical Sciences, University of Cape Town, Rondebosch, South Africa; Department of Biology, Stanford University, Stanford, CA, USA; National Genomics Center for Wildlife and Fish Conservation, Missoula, Montana, USA

**Keywords:** conservation detection dogs, faecal DNA, microsatellites, sex-biased dispersal, kinship, small carnivore

## Abstract

Advances in molecular techniques for non-invasive genetic sampling have strengthened wildlife genetics, enabling researchers to study elusive and difficult-to-study species with minimal disturbance, and conservation detection dogs can greatly increase sample numbers and the area covered. However, this combination is rarely used in Africa and has perhaps never been applied to a small carnivore on the continent. Here we use dog-assisted scat surveys to carry out the first fine-scale population genetic study of the black-footed cat (*Felis nigripes*), Africa’s smallest and most range-restricted endemic felid, and a threatened species for which conventional survey methods are largely ineffective. Using 130 successfully genotyped scat samples (nine microsatellite loci) from two populations about 650 km apart, we identified 65 individuals and assessed genetic diversity and fine-scale relatedness to explore aspects of dispersal and reproductive ecology. Both populations exhibited similar observed heterozygosity (Ho = 0.67–0.69) and allelic richness, with no heterozygote deficit (*F_IS_* = −0.02 to −0.08). Differentiation between the southern Namibia and central South Africa sites was low but detectable (*F_ST_*= 0.03–0.05). We used pairwise relatedness and candidate pedigree results to explore hypotheses about sex-specific relatedness patterns. Female–female dyads were more related than male–male dyads at the shortest distance class (0–2 km), a pattern compatible with female philopatry. There were more half-sibling than full-sibling assignments and uneven representation of candidate fathers, which may indicate polygamy and reproductive skew. These results provide baseline genetic data for a Vulnerable species and show that this approach is feasible for Africa’s small, elusive carnivores, whose dispersal, mating systems and population structure remain largely unknown. Broader spatial and temporal sampling could help assess population connectivity and reveal how threats and changing land use impact genetic diversity and structure.

## Introduction

Maintaining genetic diversity across a species’ range is a critical priority amid accelerating biodiversity loss, as it underpins population adaptive potential in changing environments and provides a key framework for linking demographic and behavioural processes to extinction vulnerability (Frankham et al., 2011; Hoban et al., 2021). For species of conservation concern, genetic metrics such as heterozygosity, allelic richness, and effective population size (*N*e) can help characterise genetic variation, demographic history, and vulnerability to future loss of diversity (Andrello et al., 2022; Sherwin et al., 2017). Crucially, *N*_e_ is shaped not only by census population size but also by population history and behavioural traits, including mating systems, dispersal patterns, and levels of reproductive skew (Anthony & Blumstein, 2000; Falconer, 1989). Polygamous mating systems can reduce *N*_e_ by increasing variance in male reproductive success, potentially accelerating inbreeding and allele loss through genetic drift (Snook et al., 2009), while sex-biased dispersal influences spatial genetic patterns (Greenwood, 1980; Pusey, 1987; Prugnolle & de Meeûs, 2002; Lawson Handley & Perrin, 2007). Advances in molecular methods applied to non-invasively collected samples have made reconstructing these processes in wild populations increasingly tractable (Waits et al., 2001; Beja-Pereira et al., 2009; Sarre & Georges, 2009).

Faecal DNA is now widely used for monitoring low-density carnivores, enabling individual identification, estimates of abundance, relatedness and genetic diversity, and assessment of fine-scale spatial genetic structure without physical capture (Long et al., 2012; Rodgers & Janečka, 2013; de Oliveira et al., 2026; Ramírez-García et al., 2025). Beyond host genetic signatures, scat can provide dietary, parasite, and microbiome information through DNA metabarcoding (Thuo et al., 2019) and can strengthen species identification where morphology is unreliable (Virtuoso et al., 2026). However, gathering sufficient samples across large areas within logistical timeframes remains a key challenge, particularly for secretive, low-density species in open landscapes. Conservation detection dogs offer a proven solution: trained to locate scats by scent, they increase sample yield and may improve spatial coverage and reduce some forms of observer or trail-based sampling bias. They also substantially improve target species discrimination in communities with multiple sympatric carnivores (Wasser et al., 2004; Reed et al., 2011; Grimm-Seyfarth et al., 2021). Detection dog deployment has been documented for more than 480 species across 62 countries (Grimm-Seyfarth et al., 2021) and has supported studies of elusive carnivores including bobcats (*Lynx rufus;* Long et al., 2007), fishers (*Pekania pennanti*; Long et al., 2007), bush dogs (*Speothos venaticus*; DeMatteo et al., 2009), and Neotropical felids (DeMatteo et al., 2014). Nevertheless, a recent global review of 338 faecal DNA studies in wildlife research reported fewer than 2% originating from Africa, and those were limited to large felids and canids (Forbes et al., 2025; Hofmann et al., 2025; de Oliveira et al., 2026). To our knowledge, the combined approach of dog-assisted scat surveys together with population genetic analyses has not been applied to small carnivores on the continent.

Small carnivores are important ecological drivers across the ecosystems they inhabit, particularly where large predators are absent or their ecological influence is limited (Do Linh San et al., 2013; Roemer et al., 2009). In much of southern Africa, large carnivores are confined to fenced reserves, elevating the importance of smaller carnivores in ecosystem regulation and trophic balance (Do Linh San, 2022). Despite their ecological value, few species have reliable population estimates, demographic data, or long-term monitoring data, substantially limiting effective conservation planning (Do Linh San et al., 2013). In addition to limited funding for studying small carnivores and their tendency to occur at low densities, their behaviour and physiology can pose additional challenges for study. Standard camera traps often have reduced detection probability for small or fast-moving species (Burton et al., 2015; Anile & Devillard, 2016; Sliwa et al., 2018) and ethical tag-mass guidelines and documented impacts of collars on behaviour and welfare also restrict the use of GPS/radio-collars for the smallest carnivores (Tomkiewicz et al., 2010; Wilson et al., 2021).

The black-footed cat (*Felis nigripes*; hereafter BFC) is an exemplary small carnivore that would benefit from non-invasive genetic sampling to address critical conservation knowledge gaps. As Africa’s smallest wild felid, endemic to the arid and semi-arid regions of southern Africa, the BFC is currently listed as Vulnerable under IUCN criterion C2a(i), with fewer than 10,000 mature individuals and a suspected ongoing decline (Sliwa et al., 2026). Primary threats include habitat loss, non-selective predator control, domestic dogs, disease, prey depletion, and competition and predation by overabundant mesocarnivores such as caracal and black-backed jackal (Sliwa et al., 2026). Because of its small body size, strictly nocturnal behaviour, cryptic colouration, and avoidance of roads and trails, conventional camera-based methods often produce low detection rates, with few available records from camera-trap surveys (Sliwa et al., 2018, 2026). Most ecological information for the species comes from long-term VHF telemetry monitoring of a small number of collared individuals at a time, conducted at a few sites (Sliwa et al., 2026).

Recent whole-genome sequencing has revealed that the species has undergone repeated population bottlenecks or long-term small, declining population size over the past two million years, resulting in some of the lowest levels of genomic diversity recorded among wild felids (Yuan et al., 2024). Reduced genetic diversity can constrain adaptive potential in the face of environmental change, a particular concern for a species already threatened by habitat degradation, climate change, and disease-related mortality, including amyloidosis, a recognised cause of mortality in both captive and wild populations (Terio et al., 2008; Sliwa et al., 2022a). Despite growing concern about potential genetic vulnerability, contemporary population-genetic assessments are lacking due to both resource constraints and the absence of scalable sampling approaches.

While recent genomic work using blood and tissue samples has begun to characterise broad-scale historical demography and phylogeography in the species (Yuan et al., 2024; Grant et al., 2026), fine-scale contemporary population structure and the behavioural processes associated with genetic diversity in wild individuals remain unknown. To address these gaps, our overarching aim was to evaluate whether dog-assisted non-invasive genetic sampling could establish contemporary genetic baselines and identify behavioural processes associated with genetic variation in wild BFC populations. We therefore conducted systematic dog-assisted scat surveys and genotyped and sexed individually identified samples using a validated microsatellite panel across two geographically distinct populations in South Africa and Namibia. We addressed three objectives: (a) to establish baseline measures of genetic diversity within and between populations, (b) to test for genetic signatures consistent with female philopatry and male-biased dispersal, and (c) to evaluate fine-scale relatedness patterns through kinship and parentage analyses to generate hypotheses about the mating system. This study provides a proof of concept that dog-led non-invasive genetic approaches can provide a practical foundation for genetic monitoring and generating ecological hypotheses for Africa’s elusive small carnivores.

## Methods

### Study Sites

We conducted dog-assisted genetic surveys at two sites within the BFC range, separated by approximately 650 km (Figure 1). The first, Benfontein Nature Reserve (BNR; 110 km²; −28.82333, 24.81964), is a private protected area located 10 km south-east of Kimberley, in the Northern Cape and Free State provinces, South Africa. The reserve lies on the flat plains of the central South African plateau, at an altitude of approximately 1,180 m. The region is semi-arid, with an average annual rainfall of 450 mm, occurring during the summer months (November to March) (Richardson, 1985). Arid plant communities dominate the vegetation, with elements of three major ecosystems: Kalahari thornveld, pure grassveld, and the Karoo (Acocks, 1975; Sliwa, 1993). The reserve is fully fenced to contain herds of indigenous ungulates; no large predators are present except the black-backed jackal (*Lupulella mesomelas*) and caracal (*Caracal caracal*). The second study area, referred to as GRU (230 km²; −27.79011, 18.30462), is in southern Namibia, near the town of Grünau in the Karas Region, and comprises private farmland lightly stocked with sheep (*Ovis aries*) managed in fenced camps. Habitat elements of karroid shrubs and mixed grasslands differ from BNR in that the region receives lower rainfall (80-120 mm/year, Atlas of Namibia Team, 2022) and is subject to predator-control activities targeting caracal and black-backed jackal, resulting in fewer potential predators than in BNR (Sliwa et al., 2022 unpublished report). Surveys were conducted in June–August 2021 at BNR and in 2022 at GRU, coinciding with the regional cooler, dry season, which is more favourable for DNA preservation and working-dog conditions (Becker et al., 2017; Wasser et al., 2004).

**Figure 1.**
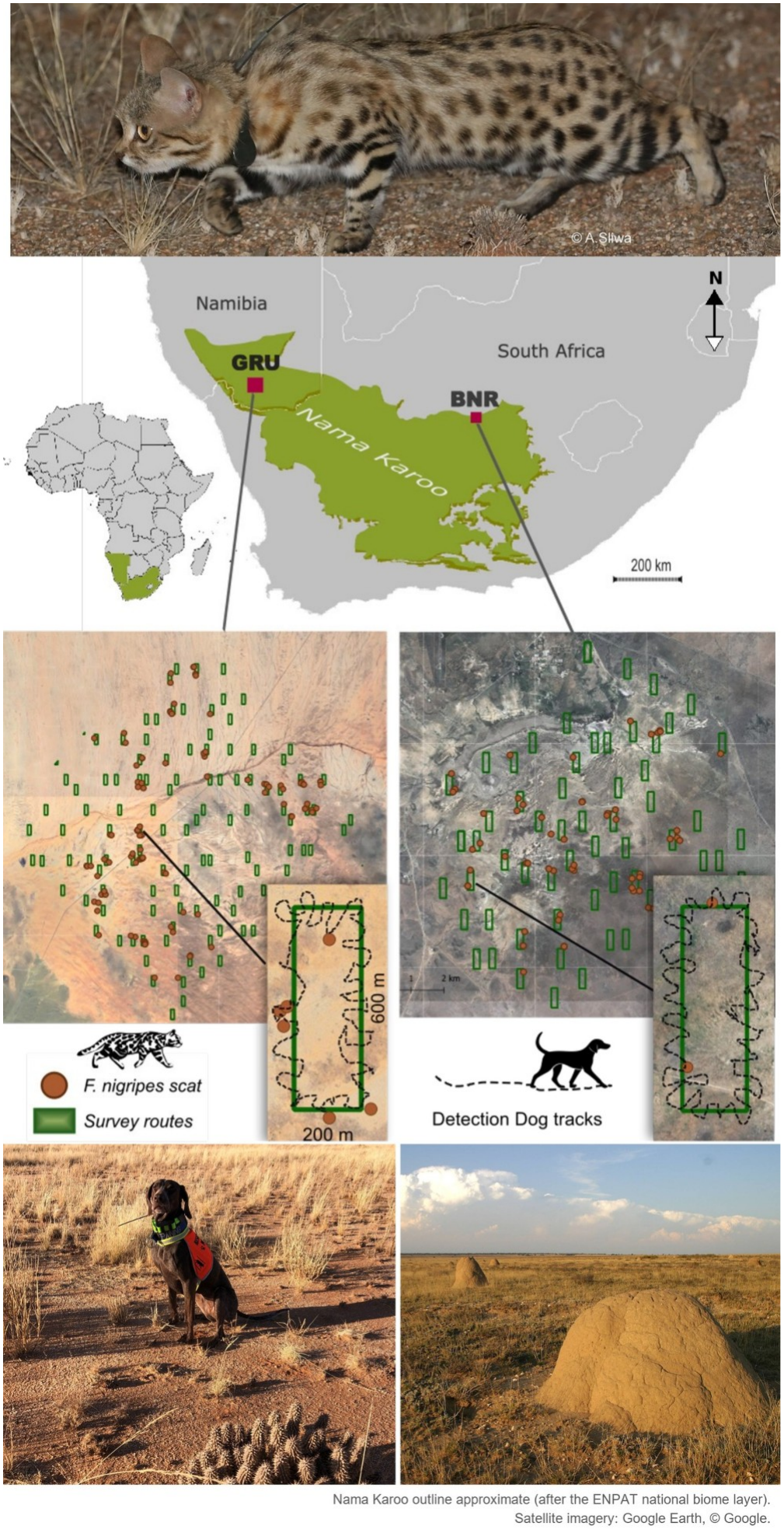
Study area and survey locations for dog-assisted scat surveys of black-footed cats (*Felis nigripes*) in southern Namibia (GRU) and the Northern Cape Province, South Africa (BNR), 2021–2022. Survey transect routes and confirmed scat locations are shown for each site. Inset panels illustrate the 200 × 600 m transect design with detection dog GPS tracks.

### Detection Dog Surveys

Because BFC scat is small and difficult to see or distinguish from co-occurring small carnivores, such as African wildcat (*Felis lybica cafra*), surveys employed a conservation detection dog with the aim of increasing sample acquisition and target-species accuracy (Long et al., 2012; Grimm-Seyfarth et al., 2021). We trained a female German short-haired pointer, ‘Lyka’, as the dedicated project dog following standard conservation detection dog training protocols (Hurt et al., 2009; Wasser et al., 2004; DeMatteo et al., 2019). The training included a 12-week program combining basic obedience, impulse control, and progressive scent imprinting using BFC scats from both wild-collared and captive individuals, using reward-based methods. We trained a passive “stand-and-freeze” indication to limit contamination. Discrimination trials included non-target felids (African wildcat; serval, *Leptailurus serval*; and caracal). Field training progressed from controlled areas with preplaced hides to blind searches within the study landscapes, confirming the dog’s ability to locate naturally deposited scats before survey deployment. South Africa currently has no national standards for certifying detection dog teams for ecological purposes, but Lyka received third-party certification for competency in ‘Black-footed Cat scent’ in 2022 from Braveheart BioDog Academy, South Africa. Research methods and Standard Operating Procedures were reviewed by the Northern Cape Province State Veterinarian to confirm that the canine used for this study is exempt from the Performing Animals Protection Act (No. 24 of 1935; PAPA). Surveys were conducted during daylight hours, when BFCs are typically resting in underground burrows, thereby limiting disturbance (Brindley et al., 2025).

We used paired, randomised, predetermined 200 × 600 m transect routes (2 per 4 km² grid cell) to achieve thorough coverage at a small female home-range scale and even sampling effort within and between sites. The dog-handler team walked transects, with the dog searching off-lead up to 100 m from the handler. GPS tracks were recorded for both the handler and the dog to quantify search effort, with the dog covering 2.5 to 3 times the distance travelled by the handler. Scats indicated by the dog were visually confirmed, given a GPS waypoint and photographed (next to a metric ruler), then collected with sterile gloves, and subsampled into specimen tubes containing colour-indicating silica gel beads for DNA preservation. The remaining scat portions were left in the field to maintain potential communication cues. We recorded relevant environmental conditions at the start of each survey and noted the approximate sample age in days (based on relative moisture and intactness), the immediate substrate, and our confidence in the target species identification. Because a stable-state freezer was unavailable, we stored samples in the most temperature-controlled space accessible at the field sites. We replaced moisture-indicating beads as needed until shipment to the lab following the 2022 field surveys, thus BNR samples were stored one year longer than GRU. The research, collection and movement of biological samples was permitted under the Department of Environment and Nature Conservation, Northern Cape Province, South Africa (Permit no. FAUNA 0636-37/2021), and Namibia’s National Commission on Research, Science, & Technology (RCIV00032018), CITES Export Permit No. 0068134. USFWS CITES Exempt. TOPS permit exempt (per comm. Department of Forestry, Fisheries and the Environment). The University of Cape Town did not require ethics approval for this study, because sampling was non-invasive (scat collection only) and the detection dog was the first author’s personal animal.

### DNA Extraction, Mitochondrial 16S Sequencing, and Microsatellite Genotyping

The National Genomics Centre for Wildlife and Fish Conservation (NGC) in Montana, USA, performed DNA extractions, sequencing, and individual genotyping. Extractions were performed using the QIAGEN QIAamp DNA Stool Mini Kit (Qiagen, Valencia, CA, USA) according to the manufacturer’s instructions. Species was determined by amplifying and sequencing approximately 360 base pairs (bp) using 16S rRNA universal primers modified from Hoelzel & Green (1992) (16SrRNA-L: 5′-TTAAACGGCCGCGGTATCC-3′; 16SrRNA-R: 5′-GAATTACGCTGTTATCCCT-3′). Reactions (30 μl) contained 50–100 ng DNA, 1× reaction buffer, 2.5 mM MgCl₂, 200 μM of each dNTP, 1 μM of each primer, and 1 U AmpliTaq Gold polymerase (Life Technologies), with cycling of 94 °C for 5 min; 34 cycles of 94 °C for 1 min, 50 °C for 1 min, and 72 °C for 1.5 min; and 72 °C for 5 min. Products were purified with ExoSAP-IT (Affymetrix-USB) and Sanger sequenced at Eurofins Genomics (Louisville, KY, USA), and the sequences were viewed, aligned, and compared with reference sequences from known species in Sequencher (Gene Codes Corporation, MI, USA). The sequence shared by all black-footed cat samples was identical to published *F. nigripes* mitochondrial genomes (GenBank accessions NC_028309.1 and KP202277.1).

Confirmed *Felis nigripes* samples were further analysed using a panel of nine microsatellite loci previously evaluated for the species: Fca08, Fca23, Fca132, Fca26, Fca43, Fca58, Fca88, Fca149, and Fca126 (Mattucci et al., 2018). Loci were amplified in multiplex PCRs (10 μl reactions containing 1 μl DNA, 1× reaction buffer, 2.0 mM MgCl₂, 200 μM of each dNTP, 1 μM reverse primer, 1 μM dye-labelled forward primer, 1.5 mg/ml BSA, and 1 U Taq polymerase; 94 °C for 5 min, then 36 cycles of 94 °C for 1 min, 55 °C for 1 min, and 72 °C for 30 s) and the products were visualised on a LI-COR DNA analyzer. Sex identification was performed using felid-specific primers for the amelogenin region, which carries a Y-chromosome deletion, as recommended for non-invasive genetic samples in felids (Pilgrim et al., 2005), with the same PCR conditions as for the microsatellite loci except for an annealing temperature of 51 °C. Replicate genotyping was performed with each sample, initially PCR-amplifying it twice across the nine variable microsatellite loci to screen for allele dropout, stutter artefacts, and false alleles (DeWoody et al., 2006). To minimise genotyping error, if any locus amplified in only one of the two replicates, or failed in both, the sample was repeated twice more. If this repeat analysis confirmed a genotype, it was retained. If the genotype failed again, the sample was assigned a missing score at the locus where it failed. Samples for which amplification failed at three or more loci were discarded from further analysis. All alleles in the dataset with low frequency (<0.05) were reamplified. Microsatellite primer testing and optimisation were assessed by the NGC using five cell and five blood samples representing 10 individuals obtained from the San Diego Zoo Wildlife Alliance Frozen Zoo (https://sandiegozoowildlifealliance.org/frozen-zoo). All ten samples yielded full genotypes.

### Screens for Quality Control

After removing poor-quality samples, the genotyping laboratory (NGC) screened the genotypes for potential errors using the program DROPOUT 2.3 (McKelvey & Schwartz, 2005) and MICROCHECKER (Van Oosterhout et al., 2004). In DROPOUT, the Error Bias (EB) test was used to detect potential genotyping errors by evaluating the distribution of pairwise mismatches among individuals. Duplicate Check (DCH) test in DROPOUT was implemented to identify loci that could artificially inflate the number of unique multilocus genotypes. The laboratory’s genotype dataset (140 samples, including the 10 cell and blood controls) was further assessed in MICROCHECKER to evaluate potential population genetic signals of allele dropout, stutter-induced scoring errors, and the presence of null alleles. The probability of identity (PI) and the probability of identity among siblings (PIsib) were calculated to assess the power of the microsatellite panel in resolving individuals (Waits et al., 2001).

### Data Analysis

#### Genetic Diversity

We assessed genetic diversity within each population (BNR and GRU) using successfully genotyped scats and a panel of nine microsatellite loci. Allelic diversity (Na), observed and expected heterozygosity (Ho, He), and inbreeding coefficients (*F_IS_*) were calculated in GenAlEx v6.5 (Peakall & Smouse, 2012). Standardised allelic richness (AR) by rarefaction was conducted using hierfstat (Goudet, 2005) in R (R Core Team, 2024) to account for sample size differences. We estimated inbreeding coefficients (*F_IS_*) per locus and population using Weir & Cockerham’s (1984) method, as implemented in hierfstat version 0.5-11. We calculated polymorphic information content (PIC) from allele frequencies to assess marker informativeness. Null allele frequencies (NAF) were estimated using the Brookfield (1996) method based on observed and expected heterozygosity. We tested for deviations from Hardy –Weinberg equilibrium (HWE) in GenAlEx using chi-square tests, with significance evaluated at α = 0.05 and a Bonferroni-adjusted α of 0.0028.

To test for a recent bottleneck, we compared observed heterozygosity with that expected at mutation–drift equilibrium given the number of alleles at each locus (Cornuet & Luikart, 1996), using 5,000 coalescent simulations per locus and site under two-phase (90% single-step mutations, variance 12; 70%, variance 30) and stepwise mutation models. Significance was assessed with Wilcoxon signed-rank and mode-shift tests (Piry et al., 1999). The simulations used our own implementation, which reproduced the output of BOTTLENECK 1.2.02 for the same input. We also tested whether FIS differed from zero by permuting alleles among individuals within each site (10,000 permutations).

Effective population size (Ne) was estimated separately for each site in two ways. First, we used the sibship-assignment method in the COLONY runs described below (Wang, 2009), assuming random mating (Alpha = 0) or using Alpha estimated from the genotypes, with 95% confidence intervals from bootstrapping. Second, as an independent estimate, we used the linkage-disequilibrium method (Waples & Do, 2008) in NeEstimator v2.1 (Do et al., 2014), assuming random mating, excluding alleles at frequency below 0.02 (0.05 for BNR, given its smaller sample), with parametric and jackknife 95% confidence intervals. Both methods assume a random sample, with respect to kinship, from a single cohort. Our samples are of mixed ages and probably include parents and their offspring, so the estimates are likely biased downwards, and no correction for overlapping generations was applied; we therefore interpret them as approximate values for the sampled animals and not as long-term Ne.

#### Population Structure

We measured differentiation between BNR and GRU using several complementary statistics to address different limitations. Pairwise *F_ST_*is the conventional measure of allele-frequency divergence and allows comparison with other studies. We calculated it in GenAlEx 6.51 using Nei’s (1973) estimator and Weir & Cockerham’s (1984) θ, which corrects for small and unequal sample sizes (the latter calculated in hierfstat). Because *F_ST_* is constrained by within-population heterozygosity and can underestimate differentiation at highly polymorphic loci such as microsatellites, we also calculated Jost’s *D* (*D_est_*), which measures allelic differentiation independently of heterozygosity (Jost, 2008). We tested the significance of each estimate with 999 permutations in GenAlEx. To identify which loci contributed most to differentiation, we calculated locus-specific *F_ST_*. We also compared allele frequencies and counted private alleles in each population. To test the null hypothesis of no genetic difference between populations, we performed analysis of molecular variance (AMOVA) in R using poppr (Kamvar et al., 2014), with 9,999 random permutations. Using multiple metrics enabled us to cross-validate population structure estimates and account for differences in the data’s sensitivity to various models. We performed Principal Coordinate Analysis (PCoA) using Nei’s genetic distance matrix in GenAlEx to visualise genetic relationships between populations.

As an additional test of differentiation, we used STRUCTURE v2.3.4 (Pritchard et al., 2000) with an admixture model with correlated allele frequencies, a sampling-location prior (LOCPRIOR = 1), and no supervised population information (USEPOPINFO = 0), with a burn-in of 100,000 iterations followed by 500,000 MCMC iterations, and 10 replicate runs for each K from 1 to 4. The most likely number of genetic clusters was evaluated using the mean log-likelihood across replicates and Evanno’s ΔK (Evanno et al., 2005), calculated from the 10 replicates per K. Because ΔK can only be calculated for intermediate values of K (here K = 2 and 3), we also considered the individual ancestry coefficients (Q), including the proportion of animals with Q ≥ 0.75 and Q ≥ 0.9. Because STRUCTURE performs poorly with few sampling sites and unequal sample sizes (Puechmaille, 2016), we repeated the analysis on 10 datasets in which GRU was randomly subsampled to the BNR sample size (n = 18) and treat STRUCTURE results as supporting rather than primary evidence of differentiation.

#### Sex-biased Dispersal

Several methods can be used to infer the dispersing sex from genotyping data using pairwise relatedness values between individuals categorised by sex. The underlying assumption is that relatedness values will be lower for the dispersing sex compared to the philopatric sex, which for BFCs is presumed to be males and females respectively. We calculated the pairwise relatedness coefficient for individuals in each population separately, using site-specific allele frequencies, in CoAncestry 1.0.1.11 software (Wang, 2011), which compares performance among seven estimators using simulated multilocus genotype data and accounts for inbreeding and genotyping errors. We selected the Triadic Likelihood Estimator (TrioML) because it has the lowest variance among estimators and tends to outperform other estimators when the sample size is low, and is recommended as a general-purpose estimator when no pedigree data are available (Wang, 2011). To test the hypothesis of male-biased dispersal and female philopatry, we categorised dyads by sex: female-female (FF), male-male (MM), and intersex (FM) to compare mean pairwise relatedness (*r*) values using one-sided Welch’s t-tests. To assess whether females are more likely to have close relatives in the population than males, we categorised pairwise *r* values into inferred relationships. Based on the empirical distribution of TrioML values across both sites, we classified dyads as first-degree relatives (*r* ≥ 0.45; parent–offspring and full siblings), second-degree relatives (0.20 ≤ *r* < 0.45), or unrelated (*r* < 0.20). We chose classification thresholds using the clustering of TrioML values observed in known parent–offspring and sibling pairs derived from kinship analysis in the program COLONY (see below) (Blouin, 2003). We calculated exact confidence intervals for binomial proportions (Clopper & Pearson, 1934). Differences among FF, FM, and MM proportions were tested using two-proportion Z-tests, implemented separately for first-degree, second-degree, and unrelated dyads.

Incorporating inter-individual distance into the pairwise relatedness analysis enables detection of fine-scale patterns of sex-based relatedness, assuming differences are most apparent at shorter distances. Because multiple scat GPS points could represent the same individual, we defined inter-individual geographic distance as the mean of all pairwise distances between samples from individual A and individual B within a site. We tested the relationship between mean dyadic *r* by sex class against ecologically motivated distance classes: 0–2 km (approximately small female home-range radius and mean nearest-neighbour distances of 1.7 km), 2–4 km (adjacent, partly overlapping home ranges), 4–8 km (adjacent, non-overlapping home ranges), and 8–16 km (broader, sub-landscape scale where movements may approach dispersal distances) (Sliwa, 2004); the 8–16 km class included GRU dyads only. We tested differences between sex classes with one-sided permutation tests in which dyad sex classes were shuffled within site (Smouse & Peakall, 1999); because individuals occur in multiple dyads, *P*-values are approximate. For each dyad × distance class, we obtained 95% confidence intervals (CIs) by nonparametric bootstrapping (10,000 resamples of dyads with replacement). We evaluated the hypothesis of female philopatry and male-biased dispersal using the mean *r* for each sex-by-distance class and calculated *P*-values as the proportion of permuted means greater than or equal to the observed mean (10,000 permutations). We ran all analyses separately for each site; if patterns were similar, we pooled the two sites after within-site dyad construction for the final analysis and visualisations. Because M35 closely matched M34 (mismatching at two of six comparable loci, with scats collected 54 m apart) and may represent a duplicate genotype, we repeated the relatedness analyses without M35 as a sensitivity test; COLONY assigned M34 and M35 as full siblings (Pr = 0.997).

Sex-biased dispersal can also be inferred through intrasex comparisons of Assignment Index Correction (AIc), which measures the likelihood that an individual originated from a sampled population (Favre et al., 1997; Mossman & Waser, 1999; Goudet et al., 2002). The underlying assumption is that the dispersing sex will have more negative AIc values due to genetic mismatches with the local gene pool. We compared mean male vs female AIc values, centred within each site and pooled across sites, for individuals with complete genotypes (the GenAlEx method cannot accommodate missing data), and tested significance using the Mann-Whitney U-test as implemented in GenAlEx, calculated in R (Peakall & Smouse, 2012). We also used the test for standard population assignment to determine the proportion of individuals assigned to the correct population (BNR or GRU) and to complement population differentiation results. These tests assume that sampled individuals are adults that have already dispersed; because ages were unknown, juveniles or subadults that had not yet dispersed may be included, which would raise short-distance relatedness in both sexes and weaken, rather than create, a sex difference (see Methodological Limitations).

#### Mating Strategy and Reproductive Skew

Although a solitary species, we explored potential family groups, as telemetry monitoring of females showed recent reproduction in the GRU study area. We analysed kinship using COLONY v2.0.7.1 (Jones & Wang, 2010). COLONY employs a full likelihood pedigree framework that jointly infers parentage, sibship, and unsampled parents, accounting for genotyping error and incomplete sampling, and is more conservative for determining kinship compared to the TrioML *r* values generated by CoAncestry. For mating system parameters, we selected female and male polygamy. This setting allows paternal and maternal half-sibships within the analysed sample. The average sibship size was set to one offspring, even though the average litter size is two (Sliwa, 2013). However, kitten survival to semi-independence is low (Sliwa et al., 2022b), and a default of one reduces false sibship assignments (Jones & Wang, 2010). As a sensitivity check, we ran COLONY with the probability that candidate mothers and fathers were present in the sample set to 50%, the standard value when the proportion of parents sampled is unknown, and to 95%, because we suspected a high proportion of the population had been sampled and wanted the model to consider sampled genotypes with missing loci as candidate parents. We also examined sensitivity to the sibship-size prior (no prior, and a prior of two offspring per parent reflecting mean litter size; Sliwa, 2013) and to a parental inclusion probability of 80%, and report results from the 95% configuration with a sibship-size prior of one (Table A1). We used default settings for other parameters for which no prior relationships were known.

To evaluate reproductive skew and assess whether parentage patterns were consistent with the predicted polygamous mating system, we used parentage and sibship assignments inferred from the Best Maximum Likelihood (BestML) configuration generated by COLONY v2.0.7.1. The BestML configuration represents the most probable pedigree structure given the genotype data and specified priors, and includes both sampled and uniquely inferred unsampled parents treated as distinct individuals. To describe realised reproduction, defined here as genotyped offspring surviving to semi-independence within the study area boundaries, we examined the maternity and paternity output files, retaining only parent–offspring assignments involving sampled parents with high assignment support (posterior probability > 0.8). Because ages were unknown, every individual was included both as a potential offspring and as a candidate parent; this allows biologically impossible multigenerational loops, which occurred only among assignments below this threshold. For population-level inference, we quantified the observed number of unique contributing fathers (F_o_) and mothers (M_o_) from the BestML configuration and compared these values to the expected number of unique fathers (F_e_) and mothers (M_e_) under random mating. We generated expected values using 10,000 randomisations, in which offspring were randomly assigned to sampled candidate males or females with replacement while holding the number of offspring constant. We summarised reproductive skew using sex-specific indices (F_e_/F_o_ for fathers and M_e_/M_o_ for mothers), where values greater than one indicate fewer contributing parents than expected under random mating. We assessed statistical significance using one-sided randomisation tests comparing observed values to their respective null distributions (Nonacs, 2000). We evaluated differences in reproductive skew between sexes using a one-sided permutation test comparing the observed difference in skew indices (F_e_/F_o_ – M_e_/M_o_) to a null distribution generated by 10,000 randomisations under random mating. As an additional qualitative indicator of mating patterns, we examined the relative proportion of full-sibling versus half-sibling relationships inferred by COLONY, as a predominance of half-siblings is expected when individuals mate with multiple partners (Jones & Wang, 2010). Because samples were obtained non-invasively and the age structure was unknown, all results are interpreted as population-level patterns across potentially overlapping cohorts rather than definitive evidence of specific mating mechanisms.

## Results

### Field Sampling and Lab Outputs

The detection dog located 185 scat samples (62 BNR and 123 GRU) during the 2021–2022 winter survey period (Figure 2). Three additional BNR scats were collected opportunistically, two from direct observation of VHF-collared individuals and one found by the handler; two samples were overly degraded and excluded from DNA analysis. Of the dog-located scats, species ID could be determined for 161 (87.0%), of which 157 were *Felis nigripes* and four matched domestic cat (*F. catus*), although some may have been African wildcat (*F. lybica*) x domestic cat hybrids. Regardless, the detection dog had 97.5% known target species specificity. Genotypes were obtained from 130 *F. nigripes* scat samples, and genotyping success differed between sites: BNR 59% [33/56] and GRU 93% [94/101] (Figure 2). Sampling quantity and genotyping success did not differ by sex across sites. The ten concessioned cell/blood samples were successfully genotyped at 9/9 loci (six females, four males), which served as controls and were excluded from further analysis. Consensus multilocus genotypes identified 18 individuals from BNR (9 females, 9 males) and 47 from GRU (21 females, 25 males, 1 of unknown sex). The Mean number of scats genotyped per individual was two and did not differ by sex (F x̅ = 2.13, M x̅ = 1.91; Mann–Whitney U test, *P* = 0.189).

**Figure 2.**
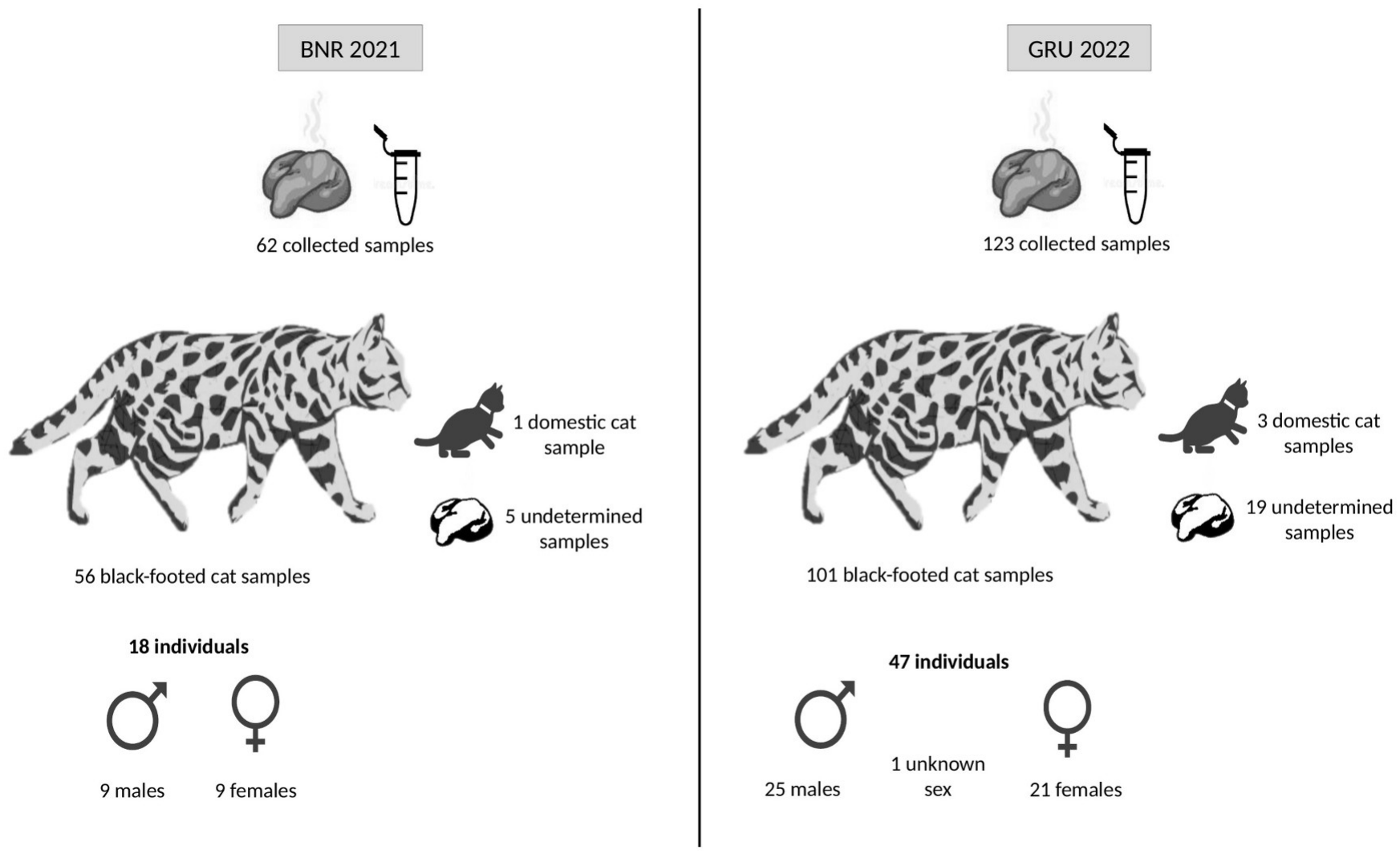
Summary of scat samples and individuals identified at Benfontein Nature Reserve (2021) and Grünau (2022). Scats located by the detection dog were identified to species by mitochondrial 16S sequencing.

### Screens for Quality Control

DROPOUT EB test results indicated that individuals differed by two or more loci, with no secondary mode detected, and did not indicate a systematic genotyping-error pattern. In the DCH test, no single locus added a disproportionate number of unique multilocus genotypes, indicating that genotyping errors were not creating false individuals, and the dataset had adequate power to distinguish individuals with a subset of five loci. The calculated PI and PIsib were 9.7 × 10⁻⁸ and 1.1 × 10⁻³, respectively, indicating the marker panel’s high discriminatory power. MICROCHECKER detected no significant homozygote excess, no evidence of stutter-induced scoring errors, no significant allele dropout, and no null alleles at any locus. Genotypes of repeat samples from the same individual (125 pairs across 32 individuals) were identical at all 1,011 locus comparisons. Marker informativeness was moderate to high, with a mean PIC of 0.60 across loci (range 0.42–0.72). Combined non-exclusion probabilities for parentage were 0.0730 (one parent known), 0.0073 (both parents known), and 0.00023 (parent pair).

Missing loci values across the 65 genotyped individuals ranged from 0 at loci Fca23 and Fca149, to 7.7% at loci Fca08, with an average of 3.2% across all nine loci. Filtering loci with >5% missing values did not significantly affect allelic diversity and resulted in slightly more negative *F_IS_* values, but the pattern was consistent across sites and did not change conclusions; thus, all loci were retained for analysis.

### Genetic Diversity

Across the nine loci, BNR had 44 alleles (mean N_a_ = 4.89 per locus), while GRU had 45 alleles (mean N_a_ = 5.00). Rarefied allelic richness was similar between populations (BNR AR = 4.85; GRU AR = 4.54). Observed and expected heterozygosity were similar between BNR (H_o_ = 0.67, H_e_ = 0.63) and GRU (H_o_ = 0.69, H_e_ = 0.63). At the population level, neither site showed evidence of inbreeding across the nine loci (*F_IS_* BNR = −0.02; GRU = −0.08).

Null allele frequencies were low, with all NAF estimates below 0.12. HWE chi-square tests identified several loci with *P*-values < 0.05; however, only GRU–Fca058 remained significant after Bonferroni correction (adjusted α = 0.0028). All other loci conformed to HWE expectations (Table A2).

Tests of heterozygosity excess did not provide evidence of a recent bottleneck at either site. Under the two-phase and stepwise models the Wilcoxon test was not significant (BNR: *P* = 0.33) under the two two-phase settings and 0.59 under the stepwise model; GRU: *P* = 0.37 and 0.82), and allele frequencies showed a normal L-shaped distribution at both sites. *F_IS_* (calculated as 1 − Ho/He, which gives slightly different values from those above) was not significantly below zero at BNR (*F_IS_* = −0.035; permutation *P* = 0.45) but was at GRU (*F_IS_* = −0.087; *P* = 0.004).

Ne estimates from the COLONY sibship method were Ne = 19 (95% CI: 10–42) for BNR and Ne = 24 (95% CI: 14–46) for GRU assuming random mating, and Ne = 26 (95% CI: 13–54) and Ne = 28 (95% CI: 17–50), respectively, assuming non-random mating. The linkage-disequilibrium estimate for GRU was Ne = 32 (95% CI: 21–52), consistent with the COLONY estimates, whereas for BNR it was unbounded (lower 95% limit = 31), rendering it uninformative.

### Population Structure

Genetic differentiation between BNR and GRU was low but statistically significant. The mean *F_ST_* across loci was 0.034 using Nei’s method in GenAlEx, while Weir and Cockerham’s estimator yielded 0.049; Jost’s Dest was 0.089 (all *P* = 0.001; Table 1). Locus-specific *F_ST_* (Nei) values ranged from 0.009 to 0.071, with the strongest differentiation at Fca058 and Fca043. A total of eight private alleles were detected in BNR and nine in GRU. Allele frequencies indicated that although many alleles were shared across populations, several loci showed notable frequency shifts. For example, at locus Fca008, allele 138 was present at 50% frequency in GRU but only 19% in BNR, and at locus Fca043, allele 125 was dominant in BNR (78%) compared to GRU (50%), although this should be interpreted with caution given 11% missing data at locus Fca008 in BNR. The AMOVA indicated significant but low genetic differentiation (Φ_PT_ = 0.095, *P* < 0.001; 9,999 permutations), with 9.5% of molecular variance attributable to differences between populations and 90.5% to within-population variation. PCoA between the two sites shows substantial overlap with shallow separation along PC1 (Figure 3). Population assignment cross-validation assigned 91% of individuals to their population of origin, with 9% assigned to the opposite site.

**Table 1.** Measures of genetic differentiation between BNR and GRU populations across nine microsatellite loci.

| **Metric** | **Value** | **P-value** |
|---|---|---|
| Fst (Nei) | 0.034 | 0.001 |
| Fst (Weir & Cockerham) | 0.049 | 0.001 |
| Jost’s Dest | 0.089 | 0.001 |
| ΦPT (AMOVA) | 0.095 | <0.001 |
| No. private alleles – BNR | 8 | – |
| No. private alleles – GRU | 9 | – |
| Mean Nm | 13.18 | – |

*Note. Fst (Nei) = Nei’s Fst calculated in GenAlEx; Fst (WC) = multilocus Weir & Cockerham’s Fst estimated in R (hierfstat); Dest = Jost’s D calculated in GenAlEx; ΦPT was calculated via AMOVA in R (poppr) on codominant genotypic distances, with missing genotypes replaced by the mean allele frequency; Nm = number of migrants per generation, calculated per locus as (1/Fst − 1)/4 from Nei’s Fst and averaged across loci. PA = number of private alleles per locus per population. Significance of Fst and Dest was assessed with 999 permutations in GenAlEx, and of ΦPT with 9,999 permutations.*

**Figure 3.**
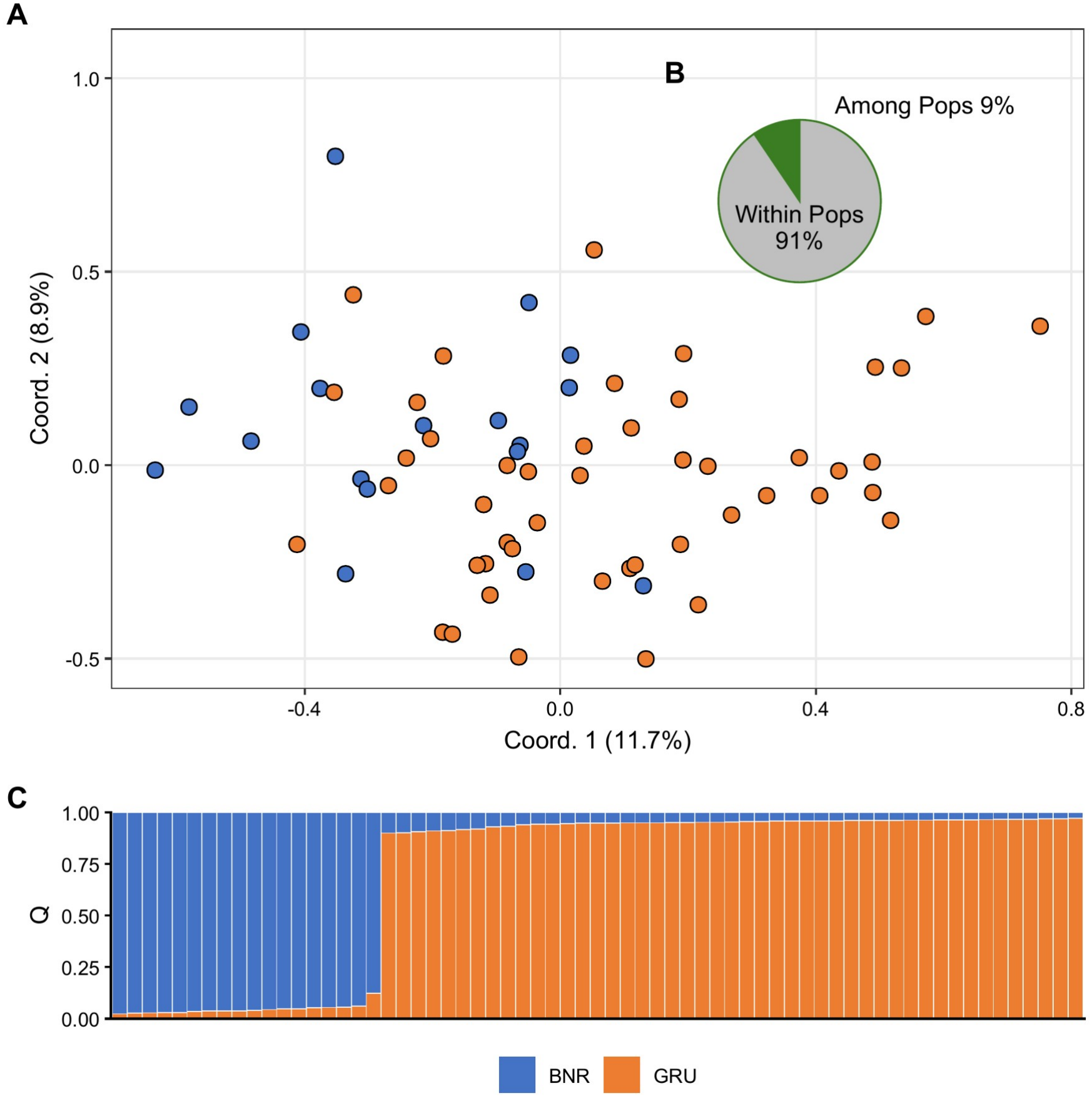
Genetic variation between BNR and GRU black-footed cat populations. (A) Principal Coordinate Analysis (PCoA) of multilocus microsatellite genotypes. (B) AMOVA variance partitioning showing among- and within-population components. (C) STRUCTURE bar plot for K = 2, with individuals ordered by sampling site.

Model-based structure analyses from STRUCTURE supported two genetic clusters corresponding to the BNR and GRU sites in the full dataset. Mean log-likelihood improved from *K* = 1 to *K* = 2 and degraded at *K* = 3–4 (−1481.53 ± 0.21 (*K* = 1), −1452.58 ± 2.06 (*K* = 2), −1614.50 ± 126.90 (*K* = 3), and −1579.03 ± 120.03 (*K* = 4)). Evanno’s Δ*K* peaked at *K* = 2 (Δ*K* = 92.70) and was minimal at *K* = 3 (Δ*K* = 1.56). BNR individuals had a mean ancestry Q = 0.955 ± 0.022 to the BNR-aligned cluster, while GRU individuals had Q = 0.949 ± 0.019 to the GRU-aligned cluster; 100% of individuals had Q ≥ 0.75, and 97% had Q ≥ 0.9 (Figure 3). However, *K* = 2 was not robust to balanced sampling: when GRU was subsampled to 18 individuals, mean log-likelihood was highest at *K* = 2 in only 2 of 10 subsets. The concordance between *K* = 2 and the differentiation metrics provides limited, descriptive evidence of site-associated allele-frequency structure (Puechmaille, 2016).

### Fine-scale Genetic Structure by Sex

#### Pairwise Relatedness

Across sites, mean pairwise *r* was higher for females (BNR F = 0.086, GRU F = 0.125) than males (BNR M = 0.051, GRU M = 0.105). The mean pairwise *r* for FF dyads (*r* = 0.120) was significantly higher than for FM dyads (0.081; t = 3.19, *P* < 0.001) and higher, though not significantly, than for MM dyads (0.099; t = 1.50, *P* = 0.067). Similarly, the proportion of dyads in the first-degree relatedness class (*r* ≥ 0.45) was highest for FF pairs (23/246 = 9.3%), intermediate for MM pairs (18/336 = 5.4%), and lowest for FM pairs (22/606 = 3.6%). Two-proportion Z-tests indicated that FF pairs had a significantly higher proportion of first-degree dyads than FM pairs (*z* = 3.38, *P* < 0.001), but FF vs MM differences were not significant. The trend was reversed when comparing the proportion of unrelated pairs among sex dyads: FF pairs had significantly fewer unrelated dyads compared to FM (*z* = −3.58, *P* < 0.001) but not MM pairs (*z* = −1.53, *P* = 0.125). Unrelated dyads (*r* < 0.20) were least common in FF pairs (183/246 = 74.4%) and most common in FM pairs (514/606 = 84.8%), with MM pairs intermediate (268/336 = 79.8%). FM pairs had marginally more unrelated dyads than MM pairs (*z* = 1.98, *P* = 0.048). These results indicate a higher fraction of close-kin pairs among females, and intersex dyads were more frequently unrelated (Figure 4). Because individuals occurred in multiple dyads, dyads are not independent, and the *P*-values of both the t-tests and the categorical comparisons are approximate; we therefore emphasise the size and direction of differences in *r* and treat these tests as exploratory (Taylor, 2015).

**Figure 4.**
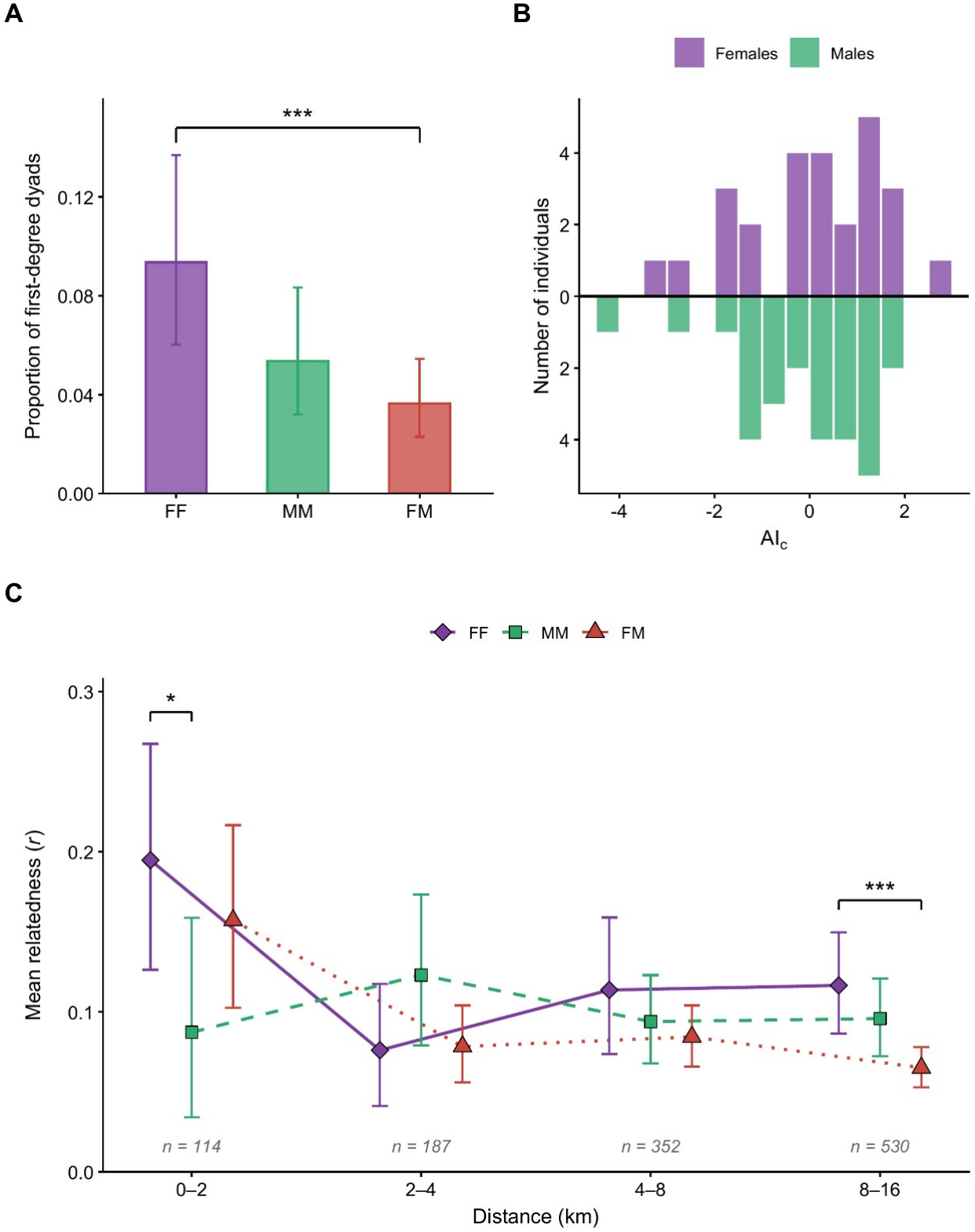
Fine-scale genetic structure by sex in black-footed cats. (A) Proportion of female-female (FF), male-male (MM), and female-male (FM) dyads assigned to first-degree relatedness (pairwise relatedness *r* ≥ 0.45; parent–offspring and full-sibling dyads), estimated using the Triadic Likelihood Estimator (TrioML) separately for each site. Error bars show Clopper–Pearson 95% confidence intervals; significant differences between dyads (two-proportion z-test) are denoted with asterisks. (B) Frequency distributions of Assignment Index Correction (AIc) values for females (above the axis) and males (below the axis) pooled across study sites (individuals with complete genotypes), with more negative AIc values indicating genetic mismatches with the local gene pool consistent with immigration. (C) Mean pairwise *r* among FF, MM, and FM dyads by geographic distance class, pooled across BNR and GRU; n is the number of dyads per class. Mean pairwise distance per dyad calculated from all genotyped scat sample locations. Error bars show bootstrapped 95% confidence intervals; asterisks denote one-sided permutation tests of FF > MM or FF > FM. \**P* < 0.05, \*\**P* < 0.01, \*\*\**P* < 0.001.

Across both sites males exhibited a lower mean AIc (−0.092 ± 0.269 SE) than females (0.095 ± 0.298 SE), although this was not statistically significant (Mann–Whitney z = 0.55, *P* = 0.581). The frequency distributions of males and females overlapped broadly, with the most negative value in males (Figure 4). These trends are directionally consistent with male-biased dispersal but lack statistical significance for firm conclusions.

#### Spatial-genetic Relatedness

At 0–2 km, FF dyads were more related than MM dyads (difference in mean *r*, Δ = 0.108; one-sided permutation *P* = 0.015), whereas FF vs FM was not significant (Δ = 0.038; *P* = 0.317). At 2–4 km, FF dyads were less related than both MM (Δ = −0.047; *P* = 0.861) and FM dyads (Δ = −0.002; *P* = 0.431). At 4–8 km, FF was not significantly higher than FM (Δ = 0.029; *P* = 0.090) or MM (Δ = 0.020; *P* = 0.211). At 8–16 km, FF exceeded FM (Δ = 0.051; *P* < 0.001) but not MM (Δ = 0.021; *P* = 0.149) (Figure 4). Excluding M35 strengthened the 0 –2 km FF vs MM difference (Δ = 0.130; *P* = 0.003) and did not change any other result (Table A3).

### Kinship and Parentage Analysis

The kinship assignment for BNR indicated that none of the 18 individuals were confidently assigned as parent-offspring pairs or full siblings; apart from four low-support links to one sampled female (F23; Pr = 0.58), individuals were assigned to unsampled parents (8 inferred mothers, 9 inferred sires). At GRU, 19 individuals were provisionally assigned as candidate offspring of four sampled parents: two sires, M53 and M31, with eight and four candidate offspring links, respectively, and two dams, F46 and F54, with six and three candidate offspring links. Two candidate offspring had both parents sampled: F40 (M31 and F46) and M44 (M53 and F46) (Figure 5; Table A4). The remaining 28 individuals were not confidently assigned a sampled parent (in total, 14 unsampled mothers and 14 unsampled sires were inferred). Full sibships were confidently assigned (Pr > 0.9) to only two pairs, while 56 pairs were identified as half-siblings (Pr > 0.9; Table A4). Assignments were sensitive to model settings (Table A1): no sampled parents were assigned when the parental inclusion probability was 50%, and no sampled sires without a sibship-size prior. F46 was assigned as a dam in all configurations with inclusion ≥ 80%, M31 and M53 as sires in all such configurations with a sibship-size prior, and M34 and M35 were full siblings in every configuration; half-sibling pairs outnumbered full-sibling pairs in all configurations except at 50% inclusion. At BNR, no parent–offspring or full-sibling relationships were confidently assigned under any setting. Because ages were unknown, the directions of parent–offspring links and individual reproductive success are treated as provisional pedigree hypotheses rather than confirmed maternity or paternity (Jones & Wang, 2010; Taylor, 2015).

**Figure 5.**
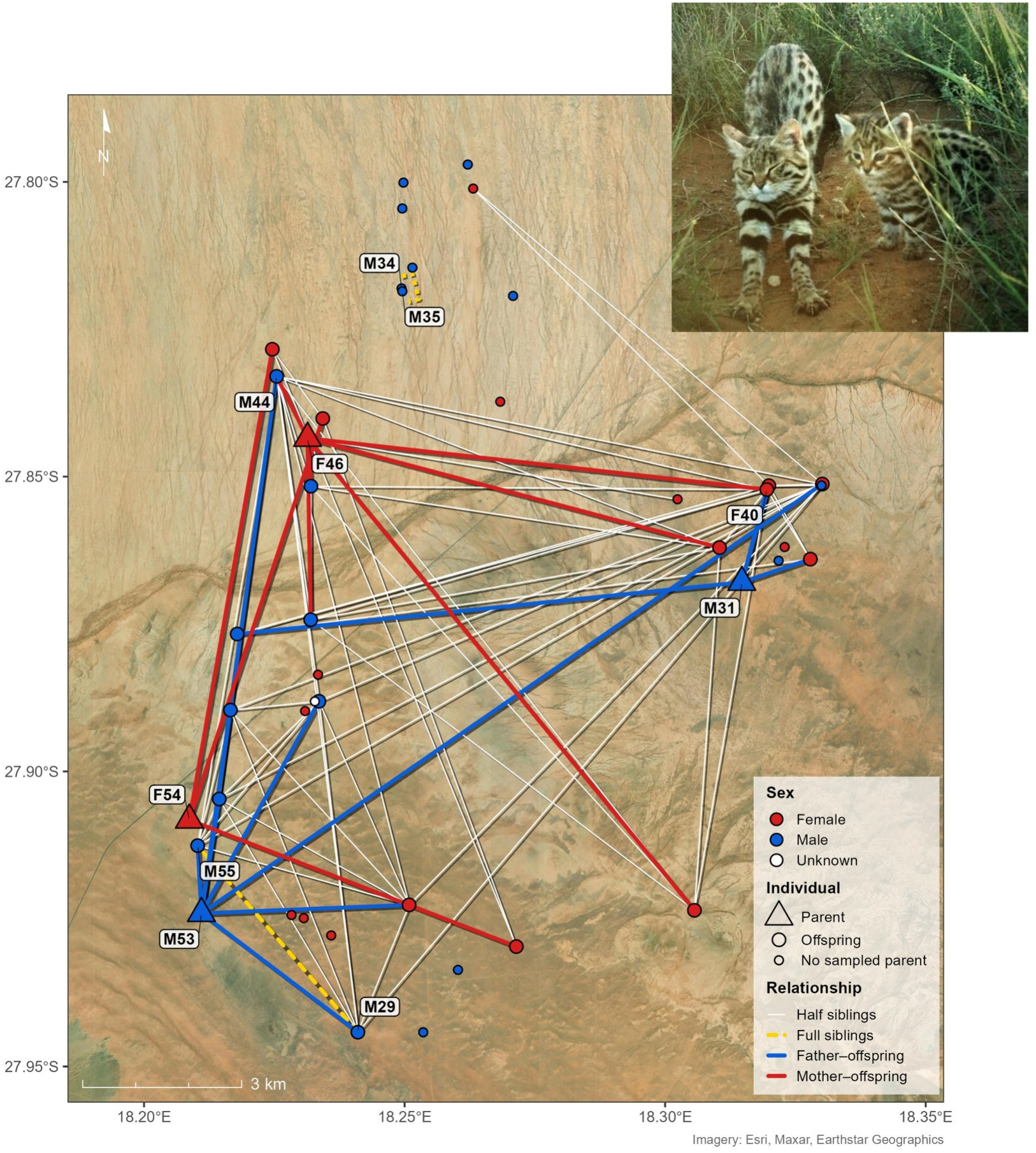
Kin relationships inferred by COLONY among black-footed cats at the GRU study area. Triangles are candidate parents and large circles their candidate offspring (posterior probability > 0.8); small circles are individuals with no sampled parent assigned. Two candidate sires, M53 and M31, were linked to eight and four candidate offspring, and two candidate dams, F46 and F54, to six and three. F46 was the candidate mother of two offspring with different candidate fathers (M31 and M53). Two full-sibling pairs and 56 half-sibling pairs were assigned (Pr > 0.9). Inset photograph: Black-footed Cat Working Group camera trap.

When assessing population-level reproductive skew in GRU, the 47 offspring were assigned to 17 inferred fathers, significantly fewer than expected under random mating (F_e_/F_o_ = 1.25; *P* = 0.007). The number of inferred mothers did not differ from random expectations (M_e_/M_o_ = 1.11; *P* = 0.131). The difference in reproductive skew between the sexes was marginal (*P* = 0.059). In BNR, inference of reproductive skew was limited by small sample size and reduced resolution of parentage assignment; thus, results are reported without further interpretation. Observed numbers of fathers (F_o_ = 9) and mothers (M_o_ = 9) did not differ from random mating expectations (paternal *P* = 1.000; maternal *P* = 1.000), and the between-sex contrast likewise indicated no difference in skew (*P* = 0.397). Because these indices depend on parentage and sibship assignments, which were sensitive to model settings, we treat them as exploratory.

## Discussion

This study provides the first non-invasive assessment of genetic diversity and sex-specific relatedness in wild BFC populations, leveraging dog-assisted non-invasive genetic sampling. The marker panel and quality-control procedures produced repeatable individual genotypes with relatively little missing data. The resulting dataset was well suited to individual identification and baseline diversity estimates, while the relatedness and pedigree analyses provide more preliminary ecological hypotheses.

The detection dog provided high species-level precision among genetically identifiable samples, a significant advantage given that BFC scats are visually indistinguishable from those of common co-occurring small carnivores, such as African wildcat (*Felis lybica cafra*) and Cape fox (*Vulpes chama*). Other carnivore scat studies have found observer bias in human-only survey teams, yielding low detection rates and/or incorrect species identification in areas with sympatric carnivores (Long et al., 2007; Lonsinger et al., 2015; Soller et al., 2020). While the use of conservation detection dogs generally increases sample yield and target-species accuracy often referred to as precision in the detection-dog literature, performance varies substantially across studies with precision rates ranging from 27% to 100% (McKeague et al., 2024). The near-98% precision achieved by the dog in this study indicates that high discriminative power can be attained with appropriate training procedures, likely enhanced by direct experience with telemetry monitoring of BFCs and access to real-world field training at the Benfontein BFC long-term study site. Because we did not survey with human-only teams, we cannot quantify the dog’s advantage directly; nevertheless, given the low density of BFCs and small scats, it is highly likely to have superseded that of human scat detection methods in the study landscapes. Notably, neither study site exhibited a female-skewed sex ratio during the sampling period, in contrast to the radio-collared sample reported by Sliwa (2004) from an earlier period. Scat-based sampling detects both resident and non-resident individuals (Wasser et al., 2004; Reed et al., 2011), and population composition may vary substantially between years; our data provide an invaluable snapshot but cannot yet distinguish these explanations.

### Genetic Diversity

The two study sites showed similar diversity, with moderately high heterozygosity and moderate allelic diversity at microsatellite loci. Heterozygosity was comparable to that reported for non-threatened felids such as the jungle cat (*Felis chaus*) and puma (*Puma concolor*), and well above that of the endangered Iberian lynx (*Lynx pardinus*), whereas allelic richness was at the lower end of the range reported for wild felids (Azizan & Paradis, 2021). These cross-species comparisons are approximate, however, because microsatellite estimates depend strongly on the marker panel and the populations sampled. Repeated sampling of relatives within genetic clusters can also bias estimates, lowering allelic richness and expected heterozygosity (Schwartz & McKelvey, 2009).

At the population level, there was no evidence of inbreeding across the nine loci in either population; instead, a slight heterozygote excess was observed (*F_IS_* = −0.02 and −0.08). By themselves, *F_IS_*values from nine loci do not identify specific processes, although slightly negative values are compatible with mechanisms that reduce inbreeding, such as dispersal and outbreeding (Pusey, 1987; Prugnolle & de Meeûs, 2002). Tests of heterozygosity excess did not indicate a recent bottleneck at either site, so we cannot attribute this pattern to demographic history, although such tests have limited power with few loci (Cornuet & Luikart, 1996; Piry et al., 1999). At GRU the heterozygote excess was significant, but the sample includes many close relatives (Table A4), which can inflate an apparent excess, so it should be interpreted cautiously. The combination of relatively high heterozygosity and low allelic diversity is nevertheless consistent with a history of small or declining population size, under which rare alleles are lost more rapidly than heterozygosity (Cornuet & Luikart, 1996; Luikart et al., 1998), and with genomic evidence of long-term small population size in the species (Yuan et al., 2024; Grant et al. 2026). Microsatellite heterozygosity is also not directly comparable with genome-wide estimates because microsatellite loci are chosen for high polymorphism and mutate rapidly; therefore, the moderately high values here do not conflict with the low genome-wide diversity reported for the species (Yuan et al., 2024; Grant et al., 2026).

Such patterns could arise in populations experiencing demographic variability while retaining genetic diversity through connectivity, where dispersal facilitates genetic mixing across a broader metapopulation despite localised allele loss (Prugnolle & de Meeûs, 2002; Szewczyk et al., 2019). However, these interpretations remain tentative. Because allelic diversity is more sensitive to long-term demographic processes than heterozygosity alone, repeated sampling over time is needed to determine whether these patterns reflect historical demographic events or ongoing processes such as gene flow and population turnover (Spencer et al., 2000).

Genetic differentiation between the sites was low but statistically detectable (Wright, 1943; Slatkin, 1993). The populations retained considerable shared microsatellite variation while also showing site-associated allele-frequency differences. Because only two sites were sampled and no intervening populations were included, the data cannot distinguish contemporary gene flow from shared ancestry or other demographic processes (Slatkin, 1985; Whitlock & McCauley, 1999; Balloux & Lugon-Moulin, 2002). However, such subtle differentiation may also arise from fine-scale kin structure, particularly in species exhibiting female philopatry, where spatial clustering of related individuals can inflate measures of genetic structure at local scales (Prugnolle & de Meeûs, 2002; Peakall et al., 2003). Consequently, the observed level of differentiation may partially reflect sampling of related individuals, although this is expected to be negligible given that most sampled individuals were unrelated. Overall, these patterns are compatible with, but do not directly demonstrate, gene flow across the region, while local demographic processes and spatial genetic structure may contribute to the weak but detectable differentiation between sites.

### Sex-biased Dispersal and Kin Structure

Building on these biologically supported genetic patterns, multiple lines of evidence are consistent with female philopatry and male-biased dispersal in BFCs. In both populations, mean *r* among females was higher than among males or mixed-sex pairs, while mean AIc values were lower in males, a directionally consistent but non-significant pattern. Fine-scale distance-class analysis further showed that neighbouring females (0–2 km) were more closely related than neighbouring males, a pattern widely interpreted as evidence of sex-biased dispersal in mammals (Greenwood, 1980; Prugnolle & de Meeûs, 2002; Peakall & Smouse, 2012). Similar sex-biased relatedness patterns have been reported in other solitary felids, including bobcats (*Lynx rufus*) and ocelots (*Leopardus pardalis*), where females are more closely related than males at close distances, and dispersal is predominantly male-driven (Janečka et al., 2006, 2007; Rodgers et al., 2015). Although our data indicate that higher female relatedness was confined to the shortest distance class; at 2–4 km, FF dyads were no more related than MM or FM dyads, and at 8–16 km FF dyads exceeded FM but not MM dyads. This pattern is compatible with philopatry, particularly at the limited spatial scale and genetic resolution of this study. Comparable findings in bobcats show that kin structure can exist without strong spatial clustering, likely due to overlapping home ranges and dispersal dynamics (Janečka et al., 2006). Given the low sampling intensity (mean ≈ two samples per individual), these results likely provide only a coarse representation of individual space use, and finer-scale kin structure would require higher-resolution sampling or integration with movement data from marked individuals. Overall, these patterns are consistent with females remaining closer to natal areas while males disperse more broadly, albeit within the spatial extent sampled.

The inclusion of opposite-sex dyads revealed additional patterns. While female–female pairs were more closely related than male–male pairs at the shortest distance class, mixed-sex pairs were less related than female–female pairs at 8–16 km, a pattern compatible with reduced local mating among close relatives, although it is not diagnostic of a specific mechanism. In solitary mammals, where stable social groups are absent, inbreeding avoidance is thought to rely primarily on sex-biased dispersal and polygamous mating systems (Pusey, 1987; Greenwood, 1980). High mortality and turnover, as reported for BFCs (Sliwa et al., 2022a), may reduce the need for long-distance dispersal if breeding opportunities become locally available, allowing young males to establish territories adjacent to natal areas rather than dispersing extensively. Such dynamics may diminish the anticipated sex differences in spatial genetic structure despite underlying dispersal biases, potentially explaining higher-than-expected male relatedness at 2–4 km (Prugnolle & de Meeûs, 2002). Across sites, male–male dyads were also somewhat more related than mixed-sex dyads (*P* = 0.068), which could reflect tolerance of related males, such as juveniles or subadults remaining in or adjacent to their fathers’ ranges; however, this difference was largely attributable to one sampled sire (M53) and his candidate sons at GRU and was not evident when that family was excluded (*P* = 0.65), so it remains a hypothesis requiring individuals of known age.

Together, the genetic diversity, dispersal, and reproductive findings align with expectations for a solitary carnivore that balances the need to avoid inbreeding with the risks associated with wide-ranging movement. Evolutionarily stable patterns of dispersal represent a balance between negative forces (mortality costs) and positive forces (reduced resource competition and inbreeding avoidance) (Bengtsson, 1978; Clutton-Brock, 1989; Lawson Handley & Perrin, 2007). Short-distance dispersal by one sex is usually sufficient to avoid inbreeding, whereas long-distance dispersal may serve to escape local population crowding (Goudet et al., 2002). Due to the limited size of the study sites, we could only examine relatedness over short distances. The decline of female relatedness beyond the shortest distance class (Sliwa, 2004) is consistent with kin structure weakening across ecologically relevant distances, although it does not by itself estimate dispersal distance. A broader sample coverage might yield insights into long-distance dispersal and the effective dispersal distance via isolation-by-distance, with important implications for assessing the extinction risk of fragmented populations.

### Reproductive Skew and its Influence on Genetic Patterns

Male-biased dispersal coupled with female philopatry is a common pattern in mammals and is widely associated with polygynous mating systems (Greenwood, 1980; Pusey, 1987; Clutton-Brock, 1989). Observational study of BFCs suggests that females tend to mate with dominant males with overlapping home ranges (Sliwa, 2004). The exploratory pedigree was compatible with unequal paternal contributions: reproductive skew was significant only among males, consistent with males mating with multiple females, although incomplete sampling, unknown ages, and model assumptions prevent direct inference about reproductive dominance. The genetic results also revealed pedigree patterns that generate hypotheses about mating-system variation that observational studies would struggle to detect. The inferred pedigree showed variation in the representation of candidate male and female parents. In the highest-likelihood pedigree, F46 was linked to six candidate offspring, with candidate offspring assigned to two sampled fathers, a pattern compatible with multiple mating across reproductive events. Sequential mating by females with different males spreads genetic and reproductive risk across cohorts and may evolve in variable environments (Watson, 1991; Yasui, 2001). Our results represent apparent fitness, often measured as relative fitness (w) or reproductive output under field or natural conditions; other demographic factors, such as survival and dispersal, can obscure reproductive patterns, especially in a species with high population turnover (Kendall et al., 1997; Lukacs & Burnham, 2005). For example, individuals might have been reproductively active, but surveys could only record offspring that survived to semi-independence and stayed in the study area. Alternative hypotheses for future testing are that this female represents a single, strong matriline occupying a high-quality territory, or inherent variation at the individual level, thereby producing more surviving offspring (Snyder & Ellner, 2024; Stover et al., 2012; Kendall et al., 2011). Furthermore, because our data lack age structure due to non-invasive sampling methods, the presence of reproductively immature individuals may reduce the effective number of breeders, thereby inflating signals of reproductive skew (Wang, 2019; Prugnolle & de Meeûs, 2002).

Under the COLONY model allowing polygamy in both sexes, the sibling structure was dominated by half-siblings, a pattern compatible with either (i) one male mating with multiple females or (ii) females mating with multiple males across reproductive events; both forms of polygamy are widely documented across mammals (Greenwood, 1980; McEachern et al., 2009; King et al., 2013). The small number of confidently assigned full-sibling pairs and the presence of numerous half-siblings, even among candidate offspring sharing a parent, are compatible with females mating with different males across reproductive events; however, they do not independently establish female promiscuity because polygamy was specified in the model (Jones & Wang, 2010; Huisman, 2017; Taylor, 2015). While multiple paternity has been demonstrated in lions and in cheetahs, whose females are solitary, evidence in other solitary felids remains limited because mating behaviour is difficult to observe (Gottelli et al., 2007; Lyke et al., 2013). Because BFCs have relatively short oestrus cycles, lasting only two nights, during which a male typically guards her closely (Sliwa, 2004), this likely limits extra-pair copulations. Mature females can produce up to three litters annually (Sliwa, 2013), and mating with different males across successive litters is plausible, though only hypothetical with our pedigree, because unaged samples cannot distinguish successive litters, years, or generations.

The lack of first-degree relatives at BNR may be an artefact of the limited number of individuals sampled. However, it is consistent with recent high mortality and reduced reproductive output documented in that population (Sliwa et al., 2022a, 2022b). Alternatively, the GRU population exhibited elevated female survival and reproduction in the year leading up to and during the sampling period (Sliwa et al., 2022b), resulting in detectable kin. Despite this recent breeding, most sampled individuals were unrelated, which we hypothesise reflected high turnover and reproductive dominance facilitated by kitten mortality and movement. The number of individuals detected at GRU likely includes juveniles and non-resident animals as well as adult residents: several females with scats inside the home range of a collared resident were closely related to one another (TrioML *r* = 0.50–0.61), consistent with juveniles or subadults still in the study area, whereas others were unrelated to them. Detection probability declines with distance from home-range centres, so parents of some individuals may have occupied ranges at or beyond the edge of the study area and remained unsampled. Conversely, resident females were likely sampled, as their home ranges contained the highest scat densities; therefore, the scarcity of parent–offspring links is more consistent with low offspring survival or dispersal than with unsampled mothers. In addition, parent–offspring and full-sibling pairs share the same expected relatedness, so with unaged samples and nine loci these relationships could not always be distinguished.

### Methodological Challenges and Future Directions

While faecal genotyping success was comparable to other carnivore studies, and exceeded expectations considering few samples were classified as fresh, 31% of dog-located samples did not produce reliable genotyping data, and future efforts could benefit from the rapid advances in genomic technologies such as short amplicon SNP panels and host DNA enrichment workflows (Natesh et al., 2019; Kraus et al., 2015; Scott & Kovach, 2024; Levengood et al., 2025). The additional year of dry storage at the BNR site may have contributed to the significantly lower genotyping success compared to GRU.

Although the 9-locus microsatellite panel used in this study performed well for individual identification and provided first-order insights and broad relatedness patterns, the modest number of loci may have limited our ability to resolve fine-scale kinship patterns with high precision. With fewer loci, relatedness estimates exhibit wider confidence intervals and reduced power to discriminate among unrelated, second-degree, and more distant kinship categories (Jones & Wang, 2010; Wang, 2016; Taylor, 2015; Weinman et al., 2015). Limited loci, small sample size, and incomplete parent sampling also increase uncertainty in COLONY pedigree assignments: they may leave some true relationships unresolved or misclassified, so the first-degree relationships identified should be treated as provisional rather than as a minimum estimate of the actual kinship structure (Jones & Wang, 2010; Taylor, 2015). Another limitation inherent to most non-invasive genetic studies is the inability to reliably assign age classes to individuals from anonymous scat samples, violating most model assumptions of a single cohort. Pooling age classes can weaken spatial genetic signals of sex-biased dispersal because juveniles and subadults of the dispersing sex may still be occupying natal ranges, thereby adding noise to potential kinship patterns expected among adults, and is known to reduce the power of spatial autocorrelation analyses and kinship estimators (Double et al., 2005). Despite this limitation, female within-home-range relatedness remained detectably higher than that of males, as expected in systems with female philopatry and male-biased dispersal. Future studies could combine non-invasive genetic sampling with DNA samples from telemetry-monitored individuals of known age and with detailed reproductive and behavioural data. Only a portion of individuals can be collared, but knowing which of them had bred, and when, would allow at least some parent–offspring relationships to be assigned with greater confidence, while non-invasive sampling provides a broader representation of the population. Repeated non-invasive sampling could then provide insight into offspring survival and movement relative to maternal home ranges, allowing the hypotheses generated here to be tested directly.

Finally, both the limited sample size and the restricted spatial extent of sampling, especially in BNR, reduced statistical power and constrained our ability to infer effective dispersal distances. Sample size can strongly influence genetic diversity estimates, and BNR sampling fell below the recommended sample size threshold of at least 25–30, ideally >30–50 individuals per population for unbiased estimates of heterozygosity and allelic richness (Hale et al., 2012; Schwartz & McKelvey, 2009; Puechmaille, 2016). Spatial genetic structure is sensitive to sampling scale relative to the species’ movement ecology, and the BNR site is only a few times the average male home-range size; it is not surprising that spatial genetic structure was weak and confined to the shortest distance class. In our study, the systematic transect design and detection dog were intended to improve sampling evenness and landscape coverage, which may have helped us detect genetic patterns in anonymous scat samples despite these limitations (Long et al., 2007; Beja-Pereira et al., 2009). These constraints reduce power and may also bias relatedness, pedigree, and spatial-genetic estimates. We therefore treat the sex-specific and reproductive results as hypotheses requiring broader sampling and higher-resolution markers. The Ne estimates should be treated cautiously: they assume a random single-cohort sample, our samples probably include relatives and several age classes, and nine loci give limited precision.

### Conservation Implications for BFCs

Using dog-assisted non-invasive genetic sampling and subsequent analyses of microsatellite loci, we established baseline patterns of diversity and sex-specific relatedness, and generated hypotheses about dispersal and reproductive structure, in two BFC populations. Together, comparable site-level diversity, weak differentiation, and sex-specific relatedness patterns are consistent with the predicted pattern of female philopatry and male-biased dispersal, while pedigree reconstruction yielded testable hypotheses about reproductive structure. While these results are encouraging, they may not represent the species elsewhere in its range, as this study has only two landscape-level data points, and populations such as GRU may fluctuate with resource availability. Future studies would benefit from larger geographic sampling, multi-year datasets, and higher-resolution markers, such as SNPs, to enable finer-scale inference of gene flow and demographic processes while maintaining compatibility across laboratories. The method could be used to monitor BFC populations across their distribution and address important conservation questions, such as which landscape types favour their persistence, the impact of human-modified landscapes, and the species’ long-term viability under current and projected environmental and land-use changes. While whole-genome sequencing studies using blood and tissue samples (Yuan et al., 2024; Grant et al., 2026) have revealed deeper historical processes influencing the species’ genetic makeup, understanding contemporary conservation challenges requires representative spatial sampling. Obtaining sufficient samples can realistically only be achieved by scaling up detection dog-led, non-invasive genetic sampling and laboratory techniques optimised for degraded scat DNA. With repeated, landscape-scale genetic sampling, practitioners could establish spatial baselines of diversity and relatedness and monitor how they change over time, providing the information needed to identify subpopulations and assess threats.

Black-footed cats face numerous threats but quantifying their impact on populations is particularly difficult without effective field surveys. While conservation status and subpopulation structure can be inferred from habitat and geographic barriers in forest-dwelling small cat species such as the African golden cat (*Caracal aurata*) or mountain species such as the snow leopard (*Panthera uncia*), as a semi-arid rangeland species with relatively contiguous habitat and no significant geographic barriers, defining subpopulation structure in BFCs requires range-wide genetic sampling and monitoring. Despite the distance (approximately 650 km) between the southern Namibia and central South Africa sites, they retain substantial shared variation. Sampling intervening populations will be necessary to determine whether that pattern reflects contemporary connectivity, historical shared ancestry, or both (Slatkin, 1985; Balloux & Lugon-Moulin, 2002). Beyond BFCs, it would be interesting to apply these methods to other small, solitary carnivores for which reproductive behaviours are virtually unknown. This study establishes that monitoring the genetic diversity of Africa’s elusive small carnivores is not only achievable but essential.

## Acknowledgements

The Black-footed Cat Working Group enabled the use of established long-term telemetry study areas, access to captured individuals, field vehicle, and species expertise. Also, thanks to the Black-footed Cat Research Project Namibia, which facilitated access to the study area and permits, and coordinated the shipment of samples. Field assistance was in collaboration with the Namibia University of Science and Technology, and graduate student Shipala Ndele. Thanks to DeBeers Consolidated Mines for continued access to Benfontein Nature Reserve, and the landowners of the Grünau farms for allowing us to survey on their private property. Collaboration with Stanford University’s Schumer Lab contributed funding to cover various field costs. Field and lab work was funded through the Panthera Small Cat Action Fund. USDA Forest Service National Genomics Center for Fish and Wildlife Conservation completed all lab analyses. Multiple individuals and organisations assisted in detection dog handling and training, including Carolynn Geary, Mike Hensman of Invictus K9, Braveheart Biodog Academy, Kayla Fratt of Canine Conservationist, Vicky Hudson of Cape Nature, Rox Brummer of Green Dogs Conservation, and Rogue Detection Dogs Team. The Lory Park Zoo, Felidae Centre, and the Cat Conservation Trust provided training sample scats.

## Data Availability Statement

The data that support the findings of this study are openly available in Dryad at http://doi.org/10.5061/dryad.k98sf7mqg. Exact scat coordinates are withheld because the black-footed cat is listed as Vulnerable and the locations are on private property; they are available from the authors on reasonable request.

## Author Contributions

Michelle M. Schroeder: Conceptualization, Data curation, Formal analysis, Funding acquisition, Investigation, Methodology, Project administration, Visualization, Writing – original draft, Writing – review & editing. Alexander Sliwa: Funding acquisition, Project administration, Resources, Supervision, Writing – review & editing. Byron Weckworth: Methodology, Project administration, Resources, Supervision, Writing – review & editing. Beryl Wilson: Funding acquisition, Project administration, Resources, Writing – review & editing. Martina Küsters: Project administration, Resources, Writing – review & editing. Victoria Grant: Investigation. Molly Schumer: Funding acquisition, Writing – review & editing. Nadine Lamberski: Resources, Writing – review & editing. Greg Distiller: Supervision, Writing – review & editing. Imogene Cancellare: Investigation. Kristine Pilgrim: Investigation. Jacqueline M. Bishop: Supervision, Validation, Writing – review & editing.

