## Supplementary material for "Dog-assisted scat sampling reveals genetic diversity, structure, and relatedness in black-footed cats (*Felis nigripes*)": Supplimental Tables A1-4

**Table A1.** Sensitivity of COLONY parentage and sibship assignments for black-footed cats to the sibship-size prior and the probability that a parent is included among the candidates, at GRU and BNR. One setting was changed at a time from the main analysis. Sampled parents are those with candidate parent–offspring assignments at posterior probability > 0.8, with the number of candidate offspring in parentheses; unsampled parents are those inferred in the best maximum likelihood configuration; sibling pairs are those with assignment probability > 0.9.

| Site | Setting | Sampled sires (offspring) | Sampled dams (offspring) | Individuals with a sampled parent | Both parents sampled | Unsampled sires / dams | Full-sibling pairs | Half-sibling pairs |
|---|---|---|---|---|---|---|---|---|
| GRU | Main analysis: inclusion 95%, sibship-size prior 1 | M31 (4); M53 (8) | F46 (6); F54 (3) | 19 | F40; M44 | 14 / 14 | 2 (M29–M55; M34–M35) | 56 |
| GRU | No sibship-size prior | none | F40 (4); F46 (4) | 8 | none | 15 / 12 | 4 (F39–F46; F39–M38; F46–M38; M34–M35) | 6 |
| GRU | Sibship-size prior 2 | M31 (3); M53 (7) | F32 (3); F42 (2); F46 (6); F54 (3) | 21 | F37; F40; M44 | 10 / 10 | 2 (M29–M55; M34–M35) | 32 |
| GRU | Inclusion 80% | M31 (5); M53 (6) | F46 (4) | 13 | F40; M44 | 12 / 17 | 2 (F39–M38; M34–M35) | 25 |
| GRU | Inclusion 50% | none | none | 0 | none | 15 / 16 | 2 (F40–M31; M34–M35) | 0 |
| BNR | Main analysis: inclusion 95%, sibship-size prior 1 | none | none | 0 | none | 9 / 8 | 0 | 3 |
| BNR | No sibship-size prior | none | none | 0 | none | 9 / 10 | 0 | 0 |
| BNR | Sibship-size prior 2 | none | none | 0 | none | 8 / 8 | 0 | 1 |
| BNR | Inclusion 80% | none | none | 0 | none | 9 / 8 | 0 | 0 |

| | | | | | | | | |
|---|---|---|---|---|---|---|---|---|
| BNR | Inclusion 50% | none | none | 0 | none | 10 / 11 | 0 | 0 |

**Table A2.** Summary of within-population genetic diversity metrics for BNR and GRU across nine microsatellite loci.

| **Pop.** | **Locus** | **$N_a$** | **AR** | **$H_o$** | **$H_e$** | **$F_{is}$** | **PIC** | **NAF** | **HWE *P*** |
|---|---|---|---|---|---|---|---|---|---|
| **BNR** | Fca008 | 7 | 7.000 | 1.000 | 0.834 | -0.168 | 0.812 | -0.075 | 0.757 |
| | Fca023 | 5 | 4.999 | 0.611 | 0.639 | 0.072 | 0.605 | 0.028 | 0.143 |
| | Fca132 | 3 | 2.999 | 0.278 | 0.434 | 0.384 | 0.386 | 0.116 | 0.163 |
| | Fca026 | 5 | 4.941 | 0.706 | 0.704 | 0.028 | 0.654 | 0.011 | 0.573 |
| | Fca043 | 4 | 3.981 | 0.444 | 0.377 | -0.153 | 0.354 | -0.041 | 0.962 |
| | Fca058 | 2 | 2.000 | 0.556 | 0.500 | -0.083 | 0.375 | -0.027 | 0.637 |
| | Fca088 | 5 | 4.990 | 0.611 | 0.696 | 0.150 | 0.644 | 0.061 | 0.015 |
| | Fca149 | 6 | 5.879 | 0.889 | 0.753 | -0.153 | 0.715 | -0.064 | 0.454 |
| | Fca126 | 7 | 6.881 | 0.941 | 0.728 | -0.264 | 0.687 | -0.109 | 0.231 |
| **BNR mean** | | **4.889** | **4.852** | **0.671** | **0.629** | **-0.021** | **0.581** | **-0.011** | **0.437** |

| **Pop.** | **Locus** | **$N_a$** | **AR** | **$H_o$** | **$H_e$** | **$F_{is}$** | **PIC** | **NAF** | **HWE *P*** |
|---|---|---|---|---|---|---|---|---|---|
| **GRU** | Fca008 | 5 | 4.344 | 0.750 | 0.639 | -0.162 | 0.580 | -0.063 | 0.919 |
| | Fca023 | 6 | 5.163 | 0.723 | 0.654 | -0.096 | 0.611 | -0.038 | 0.351 |
| | Fca132 | 4 | 3.936 | 0.442 | 0.454 | 0.039 | 0.426 | 0.012 | 0.122 |
| | Fca026 | 5 | 4.910 | 0.800 | 0.738 | -0.074 | 0.694 | -0.031 | 0.923 |
| | Fca043 | 5 | 4.111 | 0.636 | 0.641 | 0.019 | 0.582 | 0.007 | 0.925 |
| | Fca058 | 5 | 4.193 | 0.435 | 0.462 | 0.070 | 0.423 | 0.022 | <0.001* |
| | Fca088 | 5 | 4.816 | 0.739 | 0.688 | -0.063 | 0.638 | -0.026 | 0.041 |
| | Fca149 | 4 | 3.882 | 0.766 | 0.652 | -0.165 | 0.594 | -0.065 | 0.441 |
| | Fca126 | 6 | 5.480 | 0.891 | 0.696 | -0.270 | 0.653 | -0.110 | 0.007 |
| **GRU mean** | | **5.000** | **4.537** | **0.687** | **0.625** | **-0.078** | **0.578** | **-0.032** | **0.466** |

| **Pop.** | **Locus** | **$N_a$** | **AR** | **$H_o$** | **$H_e$** | **$F_{is}$** | **PIC** | **NAF** | **HWE *P*** |
|---|---|---|---|---|---|---|---|---|---|
| **Overall mean** | | **4.944** | **4.695** | **0.679** | **0.627** | **-0.049** | **0.580** | **-0.022** | **0.451** |

*Note. Na = number of alleles; AR = allelic richness (rarefied to 32 genes); Ho = observed heterozygosity; He = expected heterozygosity; Fis = inbreeding coefficient (Weir & Cockerham, 1984); PIC = polymorphic information content; NAF = null allele frequency (Brookfield, 1996); HWE P = P-value of the chi-square test of Hardy–Weinberg equilibrium in GenAlEx. *Only GRU–Fca058 remained significant after Bonferroni correction (adjusted α = 0.0028).*

**Table A3.** Sex-specific pairwise relatedness of black-footed cats by inter-individual distance class at BNR and GRU. Relatedness (*r*) was estimated with the TrioML estimator separately for each site; only within-site dyads are included. (A) Number of female–female (FF), male–male (MM), and female–male (FM) dyads per distance class, with the number excluding individual M35 in parentheses. (B) Mean *r* of FF dyads compared with MM and FM dyads; Δ is the difference in mean *r* (FF minus comparison), and *P* is from a one-sided permutation test of FF > comparison (10,000 permutations of dyad sex class within site), for all individuals and excluding M35.

**(A)**

| Distance class | FF | MM | FM | Total | BNR (FF / MM / FM) | GRU (FF / MM / FM) |
|---|---|---|---|---|---|---|
| 0–2 km | 33 | 29 (26) | 52 | 114 (111) | 6 / 3 / 16 | 27 / 26 / 36 |
| 2–4 km | 41 | 50 (46) | 96 (91) | 187 (178) | 19 / 12 / 26 | 22 / 38 / 70 |
| 4–8 km | 58 | 110 (103) | 184 (179) | 352 (340) | 11 / 21 / 39 | 47 / 89 / 145 |
| 8–16 km | 114 | 143 (133) | 273 (262) | 530 (509) | 0 / 0 / 0 | 114 / 143 / 273 |

**(B)**

| Distance class | Comparison | Mean *r* FF | Mean *r* comparison | Δ | *P* | Δ excluding M35 | *P* excluding M35 |
|---|---|---|---|---|---|---|---|
| 0–2 km | FF vs MM | 0.195 | 0.087 | 0.108 | 0.015 | 0.130 | 0.003 |
| 0–2 km | FF vs FM | 0.195 | 0.157 | 0.038 | 0.317 | 0.038 | 0.313 |
| 2–4 km | FF vs MM | 0.076 | 0.123 | −0.047 | 0.861 | −0.053 | 0.887 |
| 2–4 km | FF vs FM | 0.076 | 0.078 | −0.002 | 0.431 | −0.002 | 0.438 |
| 4–8 km | FF vs MM | 0.114 | 0.094 | 0.020 | 0.211 | 0.016 | 0.273 |
| 4–8 km | FF vs FM | 0.114 | 0.084 | 0.029 | 0.090 | 0.031 | 0.077 |
| 8–16 km | FF vs MM | 0.116 | 0.096 | 0.021 | 0.149 | 0.021 | 0.151 |
| 8–16 km | FF vs FM | 0.116 | 0.065 | 0.051 | < 0.001 | 0.051 | < 0.001 |

**Table A4.** Relationships among black-footed cats at GRU, in the best maximum likelihood configuration from COLONY: candidate parent–offspring assignments involving sampled parents (posterior probability > 0.8), with the candidate parent listed first, and full-sibling and half-sibling dyads with their assignment probabilities (> 0.9). No parent–offspring or full-sibling relationships were confidently assigned at BNR.

| Relationship | Individual 1 | Individual 2 | Probability |
|---|---|---|---|
| Candidate father–offspring | M31 | F39 | 0.857 |
| Candidate father–offspring | M31 | F40 | 0.857 |
| Candidate father–offspring | M31 | F41 | 0.857 |
| Candidate father–offspring | M31 | M63 | 0.857 |
| Candidate father–offspring | M53 | F37 | 0.961 |
| Candidate father–offspring | M53 | F75 | 0.961 |
| Candidate father–offspring | M53 | M29 | 0.961 |
| Candidate father–offspring | M53 | M44 | 0.961 |
| Candidate father–offspring | M53 | M51 | 0.961 |
| Candidate father–offspring | M53 | M55 | 0.961 |
| Candidate father–offspring | M53 | M57 | 0.961 |
| Candidate father–offspring | M53 | M71 | 0.961 |
| Candidate mother–offspring | F46 | F32 | 0.971 |
| Candidate mother–offspring | F46 | F40 | 0.971 |
| Candidate mother–offspring | F46 | F70 | 0.971 |
| Candidate mother–offspring | F46 | M44 | 0.971 |
| Candidate mother–offspring | F46 | M45 | 0.971 |
| Candidate mother–offspring | F46 | M60 | 0.971 |
| Candidate mother–offspring | F54 | F43 | 1.000 |
| Candidate mother–offspring | F54 | F47 | 1.000 |
| Candidate mother–offspring | F54 | F64 | 1.000 |
| Full siblings | M29 | M55 | 0.999 |
| Full siblings | M34 | M35 | 0.997 |
| Half siblings | F32 | F40 | 0.922 |
| Half siblings | F32 | F70 | 0.922 |
| Half siblings | F32 | M44 | 0.922 |
| Half siblings | F32 | M45 | 0.914 |
| Half siblings | F32 | M60 | 0.922 |

| | | | |
|---|---|---|---|
| Half siblings | F37 | F39 | 1.000 |
| Half siblings | F37 | F69 | 1.000 |
| Half siblings | F37 | F75 | 0.989 |
| Half siblings | F37 | M29 | 0.995 |
| Half siblings | F37 | M44 | 0.996 |
| Half siblings | F37 | M51 | 0.996 |
| Half siblings | F37 | M55 | 0.996 |
| Half siblings | F37 | M57 | 0.996 |
| Half siblings | F37 | M71 | 0.996 |
| Half siblings | F39 | F40 | 1.000 |
| Half siblings | F39 | F41 | 0.959 |
| Half siblings | F39 | F69 | 1.000 |
| Half siblings | F39 | M38 | 0.999 |
| Half siblings | F39 | M63 | 1.000 |
| Half siblings | F40 | F41 | 1.000 |
| Half siblings | F40 | F70 | 1.000 |
| Half siblings | F40 | M44 | 1.000 |
| Half siblings | F40 | M45 | 0.991 |
| Half siblings | F40 | M60 | 1.000 |
| Half siblings | F40 | M63 | 1.000 |
| Half siblings | F41 | M63 | 1.000 |
| Half siblings | F43 | F47 | 1.000 |
| Half siblings | F43 | F64 | 1.000 |
| Half siblings | F46 | F62 | 1.000 |
| Half siblings | F47 | F64 | 1.000 |
| Half siblings | F70 | M44 | 1.000 |
| Half siblings | F70 | M45 | 0.991 |
| Half siblings | F70 | M60 | 1.000 |
| Half siblings | F75 | M29 | 0.986 |
| Half siblings | F75 | M44 | 0.987 |
| Half siblings | F75 | M51 | 0.987 |
| Half siblings | F75 | M55 | 0.987 |

| | | | |
|---|---|---|---|
| Half siblings | F75 | M57 | 0.987 |
| Half siblings | F75 | M71 | 0.987 |
| Half siblings | M29 | M44 | 0.999 |
| Half siblings | M29 | M51 | 0.999 |
| Half siblings | M29 | M57 | 0.999 |
| Half siblings | M29 | M71 | 0.999 |
| Half siblings | M44 | M45 | 0.991 |
| Half siblings | M44 | M51 | 1.000 |
| Half siblings | M44 | M55 | 1.000 |
| Half siblings | M44 | M57 | 1.000 |
| Half siblings | M44 | M60 | 1.000 |
| Half siblings | M44 | M71 | 1.000 |
| Half siblings | M45 | M60 | 0.991 |
| Half siblings | M51 | M55 | 1.000 |
| Half siblings | M51 | M57 | 1.000 |
| Half siblings | M51 | M71 | 1.000 |
| Half siblings | M55 | M57 | 1.000 |
| Half siblings | M55 | M71 | 1.000 |
| Half siblings | M57 | M71 | 1.000 |
